# Rethinking animal attrition in preclinical research: expressing causal mechanisms of selection bias using directed acyclic graphs

**DOI:** 10.1101/2023.10.19.560730

**Authors:** Anja Collazo, Hans-Georg Kuhn, Tobias Kurth, Marco Piccininni, Jessica L. Rohmann

**Author notes:** **Correspondence:** Jessica L. Rohmann, Center for Stroke Research Berlin Charité – Universitätsmedizin Berlin Charitéplatz 1 10117 Berlin, Germany, jessica.rohmann /at/ charite.de; Twitter handles: @anja_collazo, @JLRohmann, @tobiaskurth, @mpiccininni3.

## Abstract

Animal attrition in preclinical experiments can introduce bias in the estimation of causal treatment effects, as surviving animals may not be representative of the entire study population. This can compromise the internal validity of the study, despite randomization at the outset. Directed Acyclic Graphs (DAGs) are commonly used tools to transparently visualize assumptions about the causal structure underlying observed data. By illustrating relationships between relevant variables, DAGs enable the detection of even less intuitive biases, and can thereby inform strategies for their mitigation. In this study, we present an illustrative causal model for preclinical stroke research, in which animal attrition induces a specific type of selection bias (i.e., collider stratification bias) due to the interplay of animal welfare, initial disease severity and negative side effects of treatment. Even when the treatment had no causal effect, our simulations revealed substantial bias across different scenarios. We show how researchers can potentially mitigate this bias in the analysis phase, even when only data from surviving animals are available, if knowledge of the underlying causal process that gave rise to the data is available. Collider stratification bias should be a concern in preclinical animal model studies with severe side effects and high post-randomization attrition.

## Introduction

The central goal of preclinical research is to determine whether a given treatment has a causal effect on a specific physiological outcome in animals, and to quantify its impact. Ideally, researchers would like to observe the outcome in the same animal at the same moment under identical laboratory and environmental conditions, only altering the individual animal’s treatment status. The unobservable outcome that the animal “would have experienced” is known as the “counterfactual outcome”.^1^ However, without the ability to travel through time, making such comparisons remains a thought experiment. Since it is not possible to observe the two counterfactual outcomes of a single animal in the real world, researchers instead target an average causal effect for a group of animals and rely on the comparison between treatment and control groups of animals, which is directly observable.

Yet, to derive internally valid estimates, it is important to ensure that, aside from treatment, everything else at the group level is as similar as possible between the two groups of animals. If the groups systematically differ in terms of the (counterfactual) risk for the outcome due to different baseline characteristics, the comparison becomes invalid. In other words the groups are not “exchangeable,” and the results may be biased.^1,2^

Preclinical researchers make use of randomized experiments to ensure, on average, this fair comparison between treatment groups.^3,4^ When researchers randomize a sufficiently large number of animals into treatment groups and carry out randomization correctly, they expect the counterfactual risk for the outcome to be on average balanced between the two groups of animals. This fulfills the condition of exchangeability between interventional groups. Without proper randomization of the treatment allocation, any observed differences between the two groups of animals may not necessarily represent the causal effect of interest. Instead, they may represent a mixing of the causal effect, if one is present, and other (non-causal) associations due to common causes that may predispose certain animals to both receiving the treatment and experiencing the outcome.^1^ This mixing, known as confounding, is well described in observational epidemiology^5^ and also recognized in preclinical literature,^6–8^ although the terminology is not consistently applied.

Another less-acknowledged threat to the internal validity is selection bias. This term refers to a systematic error in the effect estimation which arises from the exclusion or inclusion of units.^1^ In the context of preclinical research, selection bias can arise, for example, when some animals initially randomized into an interventional group are ultimately excluded from the analysis because of sudden death or mandatory euthanization due to deteriorating health during the course of the study. We refer to this such losses as animal attrition.

Especially in animal models involving invasive procedures that produce substantial pain, distress, and/or impairment, animal attrition during the course of the experiment is anticipated, yet oftentimes not transparently reported.^9–11^ A review of attrition frequencies in preclinical focal cerebral ischaemia and cancer drug studies observed uneven sample sizes in 42% of all included experiments and that animal loss often exceeded 25% of all animals used when fully disclosed.^12^ Reported sample sizes in the treatment group tended to be smaller than in the control group, indicating a higher attrition frequency in the intervention groups.^12^

Implicitly, researchers often analyze preclinical data under the assumption that animal attrition occurs at random, as if the death of an animal was only related to a random event, such as a laboratory accident. However, if the attrition is not at random, and the experimental analysis is restricted to the subsample of surviving animals only, selection bias may arise.

In this work, we focus on a specific type of selection bias called collider stratification bias. This type of selection bias, unlike other types, is unique in that it can induce bias in the analysis even when the intervention has no effect on any individual.^13,14^ To put it simply, in the presence of collider stratification bias, an association is observed *even when there is actually no true effect.* While collider stratification bias has been described in detail in other research fields, it remains entirely absent from the discussion of preclinical research results, even though it likely poses a real threat to validity.

In this paper, we show how directed acyclic graphs (DAGs), a causal inference tool, can be used to transparently illustrate causal assumptions relevant to a given experimental set up and detect potential instances in which collider stratification bias can arise. We detail a practical example from the preclinical stroke research setting and present results from a simulation study in which we quantify the magnitude of this bias and its impact on the obtained effect estimates under a variety of plausible parameter constellations.

## Methods

DAGs are visual representations of an *a priori* hypothesized causal structure. They are widely used in observational epidemiological research to transparently depict the assumed causal relationships between relevant variables under study.^1,5,15^ Briefly, DAGs consist of nodes representing measured and unmeasured variables and arrows connecting the nodes. Arrows extending from a “parent” to a “child” variable express a direct cause-effect relationship from the former to the latter. No information about the strength nor form of a relationship can be directly inferred from the arrows in a DAG. Since they are acyclic, it is assumed that no variable can cause itself directly or via another variable; this would violate the temporality principle of cause preceding effect. In a complete DAG, all direct effects among the variables as well as all common causes of any pair of variables included should be depicted. Equally important is that the absence of an arrow between two nodes reflects the strong assumption of no cause-effect relationship between them for any individual.^1,5^ Increasingly implemented across research domains outside of epidemiology, DAGs have proven useful in diagnosing the presence of potential biases and guiding study design and analysis strategies in applied research,^16–20^ but only a few examples of the use of DAGs are found in the preclinical sciences.^21,22^

### Preclinical example

We used a DAG to depict how collider stratification bias can arise in the context of preclinical *in vivo* ischemic stroke research through differential animal attrition (**Fig.1**).

**Fig. 1.**
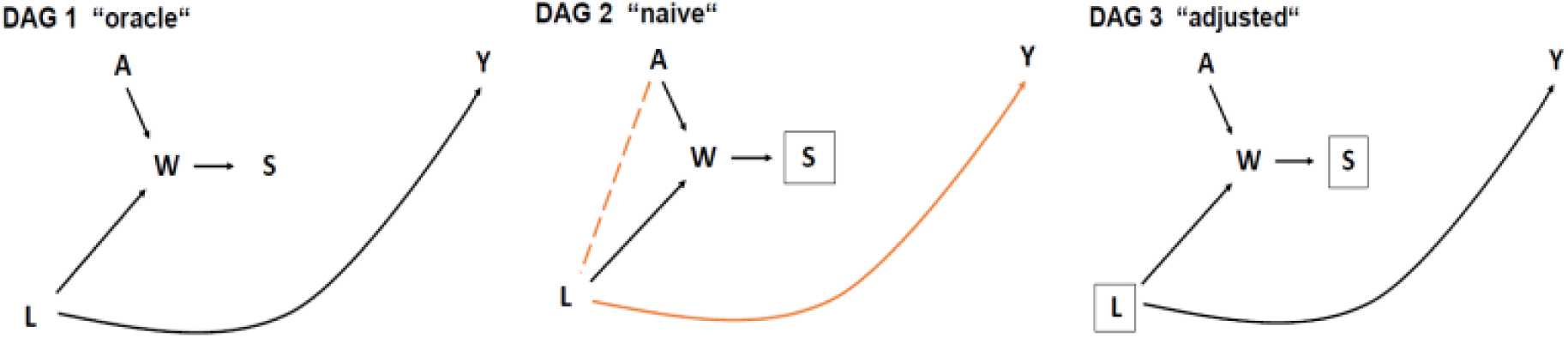
Directed Acyclic Graphs for collider stratification bias in preclinical stroke research. Variables: *A*: exposure to treatment or control; *W*: animal welfare; *S*: survival status; *L*: initial infarct volume; *Y*: final infarct volume. In this example, researchers are interested in estimating the causal effect of treatment *A* on final infarct volume *Y*. In DAG 1, hypothetically, we imagine outcome information is available irrespective of the value of survival status (oracle approach). The collider path *A* → *W* ← *L* → Y is closed (black) and the average causal effect can be estimated correctly by comparing the outcome in the two groups. In DAG 2, measuring the outcome in surviving animals only (conditioning on *S* = 1, indicated by the box around *S*) induces a spurious association between *A* and *L*, opening a non-causal path between the exposure and the outcome (orange) and will result in biasing effect estimation (naive approach). In DAG 3, adjusting for initial infarct volume (indicated by the box around *L*) closes the previously open non-causal path (now in black) and allows for the identification of the causal effect of interest (adjusted approach).

In stroke intervention experiments using an animal model, researchers often aim to derive the causal effect of a specific treatment on absolute infarct volume (in mm³) at the time point of outcome assessment. We use *A* to denote the assigned experimental group, with *A* = 1 denoting the intervention group and *A* = 0 denoting the placebo group. We use *Y* to denote the final infarct volume in mm³. The effect of interest is therefore the average effect of *A* on *Y*.

As shown in **Fig.1**, *Y* is also affected by the initial infarct volume *L,*^23^ expressed as *L* → *Y*. We define initial infarct size as an early measure of the infarct severity prior to any therapeutic intervention. The initial infarct size can vary substantially between animals depending on biological characteristics, the surgical methods used or the experimenter’s skill.^24–26^ By causing neurological deficits, inflammatory and immunological responses, which alter physiological functions, the initial infarct size *L* is also a contributing cause of the composite variable animal welfare *W*,^27,28^ which is denoted in Fig. 1 as the arrow *L* → *W.* As any treatment *A* can also have positive or negative side effects on animal welfare, we make this relationship explicit in the causal diagram with the arrow *A* → *W* (**Fig.1**). *W* is termed a collider variable on the path *A* → *W* ← *L* → *Y*, since the heads of the arrows connecting *A* → *W* and *L* → *W* “collide” into *W.*^1,13,14^ *S* represents the survival status with two possible states: *S* = 1 indicates that the animal remained in the study until the outcome was assessed. *S* = 0 indicates an unexpected loss or planned euthanization of the animal during the course of the experiment preventing the outcome assessment. The arrow *W* → *S* illustrates that the animals’ survival status, *S,* depends on their welfare status; that is, *S* is a child of *W*. Indeed, animals maintaining an acceptable welfare level throughout the experiment remain in the final analytic sample, whereas animals that drop below a certain level of welfare do not survive because their conditions substantially deteriorate and/or they must be euthanized for ethical reasons.

### Statistical analysis

For the *in silico* analyses, we translated the DAG in **Fig.1** into a set of linear structural equations under the assumption of no effect of the treatment on the outcome, with exogenous variables *A* and *L*. We considered different possible scenarios, generating the groups to have equal sizes of 5, 10 or 25 (corresponding to sample sizes, n_total_, of 10, 20 or 50).

We assumed the initial infarct size *L* to be a normally distributed continuous variable representing an absolute volumetric measure of an ischemic infarct. We set the mean and standard deviation of *L* to 25 mm³± 5 mm³, reflecting an exemplary early infarct size in a middle cerebral artery occlusion model in rats that would still be prominent enough to be detected through non-invasive imaging.^29^

Values for welfare *W* and the final infarct volume *Y* were simulated relying on the following linear structural equations:

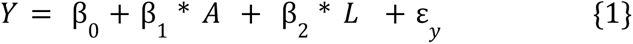

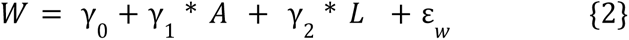

The parameter values for coefficients β_1_ (corresponding to *A → Y*), β_2_ (*L → Y*), *γ*_1_ (*A → W*) and γ_2_ (*L → W*) express the causal effects of the respective parents on the child variables in the DAG. In our simulation, we set β_1_ to zero in order to investigate the magnitude of collider stratification bias under the null (no effect of the intervention for any animal).^1^ We chose values for the other parameters in {1} in order to obtain realistic final infarct volumes. In a systematic review, O’Collins et al. reported an average absolute infarct volume of approximately 200 mm³ for an animal stroke model using Sprague-Dawley rats.^23^ Accordingly, we set β_0_ to 0, since a final infarct volume without an initial infarct volume is not conceivable. We set β_2_ to 8, which leads to an average final infarct volume of 200 mm^3^ according to {1}. The error terms *ε_y_* and *ε_w_* were assumed to be independent and normally distributed with a mean of zero and standard deviations of 10 and 2, respectively.

Since there is no standardized way of quantifying welfare scores,^28^ we generated a variable *W* with an arbitrary scale to reflect a quantified measure of welfare, with lower values indicating worse animal welfare. We set *γ*_0_ to 0 and the parameter *γ*_2_ to −1, reflecting the negative effect of initial infarct volume on animal welfare. We considered different scenarios, in which the effect of the treatment on the welfare score varied, indicating different side effect profiles of the treatment. Though often insufficiently reported, preclinical stroke studies tend to report higher animal attrition frequency for treatment groups compared to control groups, which might reflect drug toxicity.^12^ Therefore, we set *γ*_1_ to {-1, −3, −6}, reflecting potential minor, moderate, and major negative side effects of treatment.

We captured the survival of an animal throughout the experiment in our simulations using the binary variable *S* (“Survival Status”), which was deterministically obtained from *W*. In accordance with reported attrition frequencies for *in vivo* stroke research,^12^ we simulated the exclusion of animals with a welfare score below the 20th-(low attrition scenario), 30th-(moderate attrition scenario) or 50th-(high attrition scenario) percentiles of all animals in the simulated dataset. The different attrition frequencies are intended to reflect plausible differences in the severity of the animal models or environmental conditions particularly conducive (or detrimental) to survival.

Since we considered a continuous outcome, the treatment effect was estimated as the absolute difference between the mean final infarct volumes of animals in the treatment versus control groups. An effect estimate with a negative sign thus indicates a beneficial treatment effect.

We estimated the treatment effect using three approaches (in accordance with the identification strategies shown in **Fig. 1**):

1.) The “oracle” approach: the effect estimate was calculated as the difference in the mean final infarct volume between treated and untreated animals using all data points, including animals with *S* = 0. Obviously, this hypothetical, all-knowing approach is not feasible in the real world, since the variable *Y* cannot be collected for animals with *S* = 0; these animals would not have actually survived to the time point of outcome assessment.
2.) The “naive” approach: The treatment effect was estimated as the difference in the mean final infarct volume between treated and untreated groups *only* among those animals that survived until the time point of outcome measurement (restricting to *S* = 1), mirroring real-world conditions.
3.) The “adjusted” approach: The treatment effect estimate was obtained from a linear regression with final infarct volume as the dependent variable and treatment status and initial infarct volume as independent variables. The regression was fit *only* among animals that survived until the time point of outcome assessment (restricting to *S* = 1), mirroring real-world conditions. The regression coefficient for treatment thus represents the estimated effect of the treatment, adjusting for *L*.

In total, we created 27 distinct scenarios with all possible constellations of the parameters (small, medium, or large sample size; minor, moderate, or major side effects; low, moderate, or high attrition). For each scenario, the simulation was repeated 10,000 times. We calculated the mean and the 2.5th and 97.5th percentiles of the effect estimates obtained using the three aforementioned approaches (oracle, naive, adjusted) across the 10,000 runs.

We used R version 4.2.1 and RStudio version 2022.12.0 for all analyses and visualizations. The corresponding R code, figures, and tables can be retrieved from our Git repository (https://github.com/collazoa/Rethinking_animal_attrition).

## Results

Given the DAG 1 in **Fig.1**, both treatment and initial infarct volume affect welfare score, and they are independent of each other. However, the complete-case analysis (naive approach, **Fig.1**, DAG 2) involves estimating the association of interest only among the stratum of surviving animals (*S* = 1). This selection conditions on a child of the collider *W* (indicated by the box around *S* in **Fig.1**, DAG 2). The restriction to analyzing data from surviving animals only thus induces a spurious association via a collider path, as shown by the orange dashed line in **Fig.1** DAG 2.^5,13^ Thus, the measured association between *A* and *Y* in the data does not reflect the causal effect of interest, but instead a mixing between the effect of interest (should one be present) and an additional non-causal association introduced by the open path *A → W ← L → Y*.

From the assumed data generation mechanism (structural equations, distributional assumptions and chosen parameters), it follows that animals have a higher probability of dying or being euthanized during the course of the study if they were in the treatment group (*A* = 1) due to negative side effects or if they had large initial infarct volumes (*L)*. Since having received treatment and having large initial infarct volumes are both causes of low welfare scores, among the surviving animals (*S* = 1), treated animals are less likely to have large initial infarct volumes. Given that final infarct volume (*Y*) increases with the initial one (*L)*, smaller initial infarct volumes in the surviving treated animals result in smaller final infarct volumes compared with surviving control animals.

Consequently, in such scenarios, the naive approach to effect estimation (complete case analysis of surviving animals) is biased in favor of the treatment because of the treatment’s negative side effects despite the absence of any causal treatment effect (see an example in **Fig.2**).

**Fig. 2.**
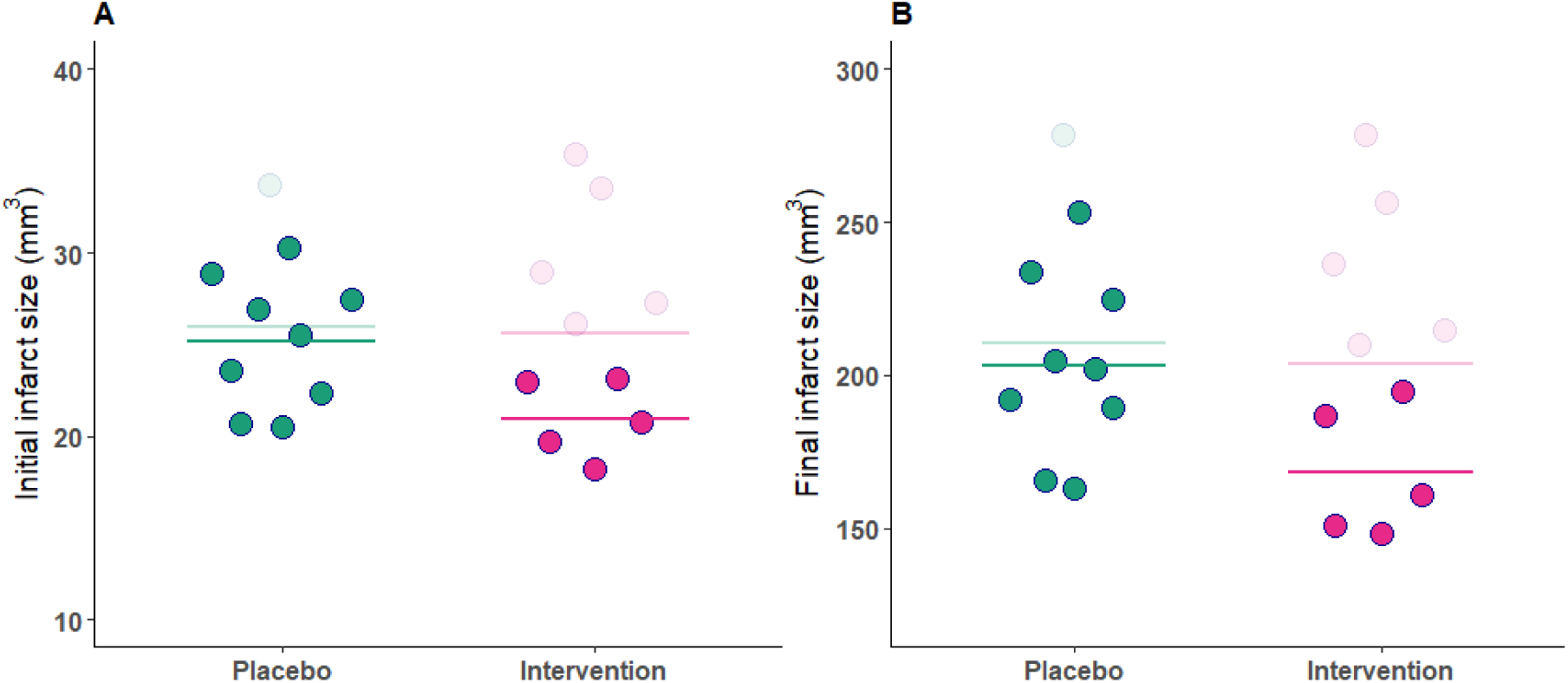
Example of impact of collider stratification bias on initial and final infarct sizes using simulated data. Values for initial infarct volume, *L*, and final infarct volume,*Y*, are shown. The solid dots and lines represent data from surviving animals (*S* = 1) and the mean values for the underlying infarct size *L* and observed outcome *Y* in surviving animals of each experimental group (naive approach). The semi-transparent data points represent (hypothetical) values of censored animals (*S* = 0). The semi-transparent lines depict the means of *L* and *Y* in the hypothetical dataset without censoring (oracle approach). **A:** At study outset, both experimental groups show similar means for the initial infarct size *L* (semi-transparent line). Due to the underlying data generation mechanism, remaining animals in the treatment group show - on average - smaller initial infarct sizes (solid points). **B**: Since *Y* increases with increasing *L*, the surviving animals of the treatment group show smaller final infarct sizes despite absence of an actual causal effect.

The results from our simulation of 10,000 experiments for each set of parameter constellations in the assumed data generation mechanism are shown in **Fig.3**.

**Fig. 3.**
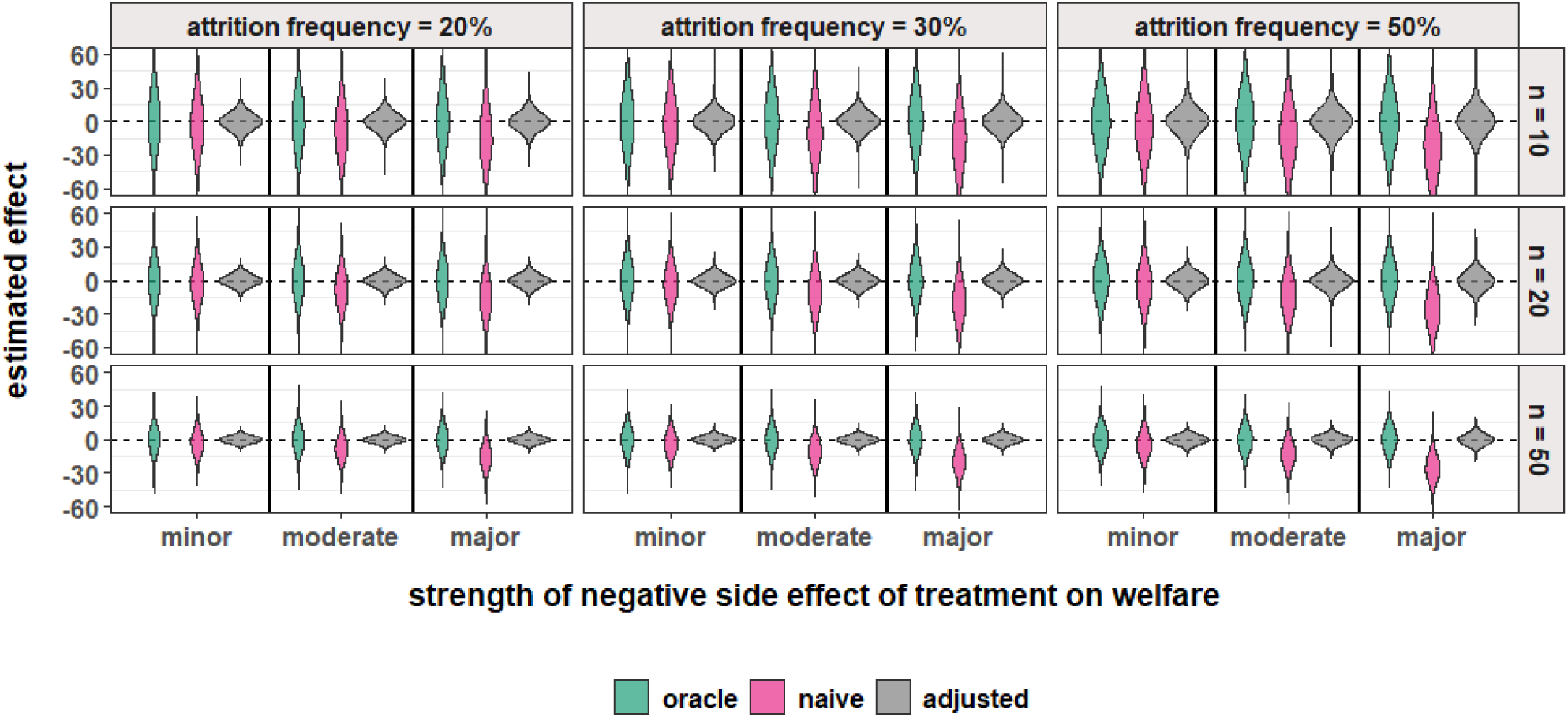
Simulation results: Three approaches for effect estimation under different attrition frequencies, strengths of side effects and sample sizes. Distribution of effect estimates for 10,000 simulated experiments for each of 27 scenarios created from the combination of different total sample sizes, n (10, 20 or 50), strength of side effects, *γ*_1_ (major, moderate, minor) and attrition frequency (20%, 30%, 50%). Results are shown separately for the oracle approach (green): complete dataset E(Y |A=1) − E(Y |A=0); the naive approach (pink): with censoring of animals according to welfare score, E(Y |A=1,S=1) − E(Y |A=0,S=1); and the adjusted approach (gray): with censoring of animals according to welfare score and adjustment for baseline infarct volume, E(Y |A=1,S=1,L=l) − E(Y |A=0,S=1,L=l). While the distributions of the oracle approach (green) and the adjusted approach (gray) are centered approximately around the true causal effect (zero), the average effect estimate obtained by the naive approach appears to deviate substantially from zero under multiple scenarios.

In 0.5% of the simulated experiments, the effect estimates for the naive and adjusted approach could not be calculated. For example, in the scenario with very few animals (n_total_=10) and high attrition (50%), these effect estimates could not be computed 10% of the time. We excluded these missing values from calculation of mean and quantiles of the effect estimates.

The mean causal effect estimate obtained using the oracle approach was approximately zero across all 27 scenarios (**Fig. 3** and **Table 1**). Indeed, as expected, this approach is unbiased. In contrast, the mean effect estimates obtained using the naive approach differed from the true causal effect (zero), indicating bias (**Fig. 3** and **Table 1**). This result shows that only including surviving animals (*S* = 1, complete-case analysis) can induce an association between the treatment and the outcome merely due to the collider stratification bias. The magnitude of this bias varied widely across the different scenarios and increased with increasing attrition frequencies and side effect severity (*γ*_1_). When the attrition rate was low (20%) and the side effects were minor, the bias for the naive approach was −2.3 mm^3^ (2.5th and 97.5th percentiles: −50.9, 47.8) when the sample size was 10 and −2.8 (−23.8, 18.5) when the sample size was 50 (Table 1). With higher attrition (50%) and major side effects, the quantified bias observed using this approach was as large as −24.5 mm^3^ (−83.2, 32.9) for a sample size of 10 and −25.7 mm^3^ (−51.0, −1.2) for n_total_=50 (**Table 1**). While improving the precision of the effect estimates (less variability), increasing the sample size did not reduce the naive approach’s bias induced by collider stratification.

**Table 1.** Effect estimates obtained using three different approaches under different scenarios of sample size, attrition frequencies, and side effects.

| Attrition frequencies (%) | Side effects, severity | Oracle approach: mean (mm<sup>3</sup>) | Oracle approach: 2.5% and 97.5% quantiles (mm<sup>3</sup>) | Naive approach: mean (mm<sup>3</sup>) | Naive approach: 2.5% and 97.5% quantiles (mm<sup>3</sup>) | Adjusted approach: mean (mm<sup>3</sup>) | Adjusted approach: 2.5% and 97.5% quantiles (mm<sup>3</sup>) |
| --- | --- | --- | --- | --- | --- | --- | --- |
| **n = 10** |  |  |  |  |  |  |  |
| 20 | minor | 0.2 | -50.7, 52.3 | -2.3 | -50.9, 47.8 | -0.1 | -15.8, 15.6 |
|  | moderate | -0.5 | -51.7, 51.0 | -8.0 | -56.6, 41.3 | -0.1 | -16.5, 16.2 |
|  | major | 0.2 | -49.7, 51.1 | -13.4 | -63.2, 37.5 | 0.0 | -16.2, 16.4 |
| 30 | minor | 0.0 | -51.3, 51.4 | -2.9 | -52.7, 46.2 | 0.0 | -17.6, 17.7 |
|  | moderate | 0.2 | -51.4, 50.5 | -9.1 | -59.9, 40.6 | 0.0 | -17.9, 17.7 |
|  | major | 0.6 | -51.0, 51.9 | -17.4 | -68.4, 34.3 | 0.1 | -18.3, 19.0 |
| 50 | minor | 0.2 | -51.0, 51.7 | -4.3 | -59.7, 50.8 | -0.1 | -25.4, 25.0 |
|  | moderate | 0.3 | -51.2, 52.0 | -12.3 | -70.3, 44.0 | -0.3 | -27.8, 25.2 |
|  | major | 0.0 | -49.2, 51.4 | -24.5 | -83.2, 32.9 | 0.0 | -29.4, 29.9 |
| **n = 20** |  |  |  |  |  |  |  |
| 20 | minor | -0.1 | -36.1, 36.7 | -2.7 | -36.4, 30.8 | 0.1 | -10.4, 10.3 |
|  | moderate | 0.1 | -36.5, 36.8 | -7.9 | -42.3, 25.8 | -0.1 | -10.8, 10.3 |
|  | major | -0.2 | -36.1, 36.5 | -14.6 | -49.1, 20.3 | 0.0 | -10.5, 10.9 |
| 30 | minor | 0.2 | -35.6, 36.1 | -3.3 | -37.1, 30.4 | -0.1 | -11.3, 11.2 |
|  | moderate | -0.3 | -36.1, 35.2 | -10.2 | -44.2, 24.6 | -0.1 | -11.5, 11.4 |
|  | major | -0.1 | -35.9, 35.9 | -19 | -55.1, 17.0 | 0.1 | -12.1, 12.4 |
| 50 | minor | 0.2 | -36.0, 36.6 | -4.3 | -42.3, 32.9 | 0.0 | -13.9, 13.8 |
|  | moderate | 0.1 | -35.7, 36.1 | -12.9 | -51.4, 25.4 | -0.1 | -15.0, 14.5 |
|  | major | 0.0 | -36.8, 35.7 | -25.4 | -67.4, 14.9 | -0.1 | -16.8, 16.8 |
| **n = 50** |  |  |  |  |  |  |  |
| 20 | minor | 0.1 | -22.8, 23.5 | -2.8 | -23.8, 18.5 | 0.0 | -6.3, 6.3 |
|  | moderate | 0.1 | -22.9, 23.1 | -8.1 | -29.3, 13.3 | 0.0 | -6.3, 6.2 |
|  | major | 0.0 | -23.4, 23.5 | -15.2 | -36.9, 6.7 | -0.1 | -6.6, 6.3 |
| 30 | minor | 0.0 | -22.9, 22.8 | -3.5 | -24.6, 17.9 | 0.0 | -6.9, 6.7 |
|  | moderate | 0.0 | -22.5, 22.8 | -10.1 | -31.5, 11.1 | -0.1 | -6.9, 6.8 |
|  | major | 0.0 | -23.1, 23.1 | -19.4 | -42.0, 3.0 | 0.0 | -7.3, 7.4 |
| 50 | minor | 0.1 | -22.8, 22.6 | -4.4 | -27.7, 18.4 | 0.0 | -8.3, 7.9 |
|  | moderate | 0.0 | -23.3, 22.8 | -12.8 | -36.4, 10.7 | 0.1 | -8.5, 8.5 |
|  | major | 0.2 | -22.6, 22.6 | -25.7 | -51.0, -1.2 | 0.1 | -9.8, 9.7 |

Mean and 2.5th and 97.5th percentiles of effect estimates for 10,000 simulated experiments for each of 27 scenarios created from the combination of different total sample sizes, $n$ (10, 20 or 50), strength of side effects, $\gamma_1$ (major, moderate, minor) and attrition frequency (20%, 30%, 50%). Results are shown separately for the oracle approach: complete dataset $E(Y | A=1) - E(Y | A=0)$ ; the naive approach: with censoring of animals according to welfare score, $E(Y | A=1, S=1) - E(Y | A=0, S=1)$ ; and the adjusted approach: with censoring of animals according to welfare score and adjustment for baseline infarct volume, $E(Y | A=1, S=1, L=I) - E(Y | A=0, S=1, L=I)$ .

We note that while attrition frequencies were similar for both groups in the presence of minor side effects, this is not the case when moderate or large side effects were present. As expected, according to our data generation mechanism, severe side effects associated with the treatment lead to higher attrition rates in the treatment group (**Table 2**).

**Table 2.** Attrition frequencies stratified by experimental group.

| Side effects, severity | Total attrition frequencies (%) | Attrition frequency in treatment group (%) | Attrition frequency in control group (%) |
| --- | --- | --- | --- |
| minor | 20<br>30<br>50 | 23<br>33<br>54 | 17<br>27<br>46 |
| moderate | 20<br>30<br>50 | 28<br>40<br>62 | 12<br>20<br>38 |
| major | 20<br>30<br>50 | 35<br>49<br>73 | 5<br>11<br>27 |

*Presentation of the attrition frequencies by experimental group under different parameter values for negative side effects $\gamma_1$ ( $A \rightarrow W$ ) (minor, moderate, or major) and total attrition frequencies, (20%, 30% or 50%).*

When using the adjusted approach, we condition on *L*, blocking the non-causal path opened by the selection on S (*A → W ← L → Y*). Therefore, this approach provides unbiased estimates for the causal effect conditional on *L*.

In our data generation mechanism, this conditional effect is equivalent to the marginal (i.e., average) one, which was targeted in the oracle approach. For this reason, the adjusted approach yielded mean effect estimates close to zero across all 27 scenarios (**Table 1**). Since the adjusted approach targets a conditional causal effect rather than a marginal one, not only did the adjusted approach lead to unbiased estimation of the true effect but was also more precise. For example, in the scenario with large sample size, high attrition and major side effects, using the oracle approach, we obtained a mean effect estimate of 0.2 (−22.6, 22.6), while using the adjusted approach, a mean estimate of 0.1 (−9.8, 9.7).

## Discussion

In this work, we used DAGs to illustrate how differential animal attrition can induce collider stratification bias, a type of selection bias, in the estimation of causal effects in preclinical disease models. We present an example from preclinical stroke research in which the animal’s welfare is affected both by the treatment (i.e., negative side effects) and the initial infarct volume. Since the outcome, final infarct volume, is affected by the initial infarct size, we show how collider stratification bias can lead experimenters to erroneously conclude that a treatment has a beneficial effect even when no real effect exists.

The bias in this practical example arises from the fact that analyzing data only from the subset of animals that survived (“naive approach”) corresponds to implicitly conditioning on a (child of a) collider variable, thereby inducing a non-causal association between the treatment and the initial infarct size. We quantified the magnitude of the collider stratification bias in a simple model and showed that, being a systematic error, this bias cannot be reduced by increasing the sample size.

We illustrate that researchers can potentially mitigate this bias in the analysis phase, even when only data from surviving animals are available, by measuring and statistically adjusting for variable(s) on the open non-causal path (“adjusted approach”). This requires knowledge about the underlying causal structure that gave rise to the data, for which a DAG can be a useful visualization tool. In our simple example, by measuring the initial infarct size and including it as a covariate in the regression model fit using the data of the surviving animals, we could completely eliminate the collider stratification bias even under high animal attrition and severe side effects.

Selection bias, though less intuitive than measurement error or confounding, has been described in the context of preclinical (stroke) research.^12,30–32^ We show how the use of a causal framework, specifically, through use of DAGs, can alert us to the presence of these biases and other threats to internal validity and inform the analysis in a transparent way. Collider stratification bias, unlike other selection biases, can lead to misleading effect estimates even when no effect is actually present.^1^ Therefore, it is plausible that erroneous decisions regarding the progression of treatments to confirmatory preclinical trials or first-in-human studies could be based on this bias, raising ethical concerns with regards to animal use and futile testing in patients.^33,34^ While we focus on the specific case in which the treatment has no effect in any animal here, we emphasize that collider stratification can also induce bias by the same mechanism when the treatment has an effect on the outcome.^1^

### Limitations

While DAGs represent non-parametric models and the nature of collider stratification bias is agnostic to the specific type of functional relationship, we made several simplifying assumptions in our application.

First, we included a limited number of variables in our DAG, to avoid overcomplication and focus on the simplest structure of collider stratification bias. In practice, for example, the presence of other common causes of animal welfare and final infarct volume such as infarct location should also be considered and depicted in the DAG. In general, infarct volumes in preclinical stroke models differ by the methodological approach employed; the stroke does not arise on its own but is induced in the laboratory using techniques such as transient or permanent intraluminal thread occlusion middle cerebral artery occlusion (MCAo).^23,35^ These models are heavily dependent on operational or environmental factors such as type of anesthesia, duration of ischemia, or surgical skill.^24,35^

Second, we assumed the structural causal equations for animal welfare and the final infarct volume were linear, and the treatment had no effect on the outcome. Due to these assumptions, the conditional causal effect estimated by the adjusted approach also represented an unbiased estimate of the marginal causal effect, which is typically targeted in randomized controlled experiments. This is also the reason why the adjusted approach in this specific data generating mechanism yielded more precise estimates compared with the oracle approach.

These assumptions likely represent an oversimplification. In the real world, it is plausible that the treatment has an effect on the outcome that varies across animals. In the presence of such effect measure modification (or if the interest lies in a non-collapsible effect measure), more complex statistical methods, such as g-methods,^1^ should be employed to adjust for the selection bias.

Third, we assumed independent errors in our structural causal model. We recognize that this cross-world assumption is untestable.^1^

Fourth, the magnitude of the collider stratification bias depends on the parameters in the structural causal model. For example, the initial infarct volume must have a strong impact on the final infarct volume in order for the collider stratification bias to be non-negligible in the naive approach. Parameter choices as well as the assumed linearity of cause-effect relationships in the DAG limit the generalizability of the bias estimates that we have reported.

Fifth, we acknowledge that to adjust for the selection bias during the analysis phase, it is necessary to correctly model the relationship between the variables. In our didactic example, we knew that the model was correctly specified. However, whether a model is correctly specified is not generally known nor testable. Using a model that misspecifies the functional relationships between variables can introduce bias. This problem presents a substantial challenge in preclinical research considering the typically low sample sizes.

Lastly, we opted to exclude the small number of effect estimates in the naive and adjusted approaches that could not be computed due to too few animals in one particular stratum. We acknowledge that excluding these missing estimates may have introduced some bias in the calculated means and quantile ranges.

### Implications

Our results emphasize the need to strive for low animal attrition during the course of an experiment to maintain internal validity in preclinical research. Since higher survival probabilities of animals can improve internal validity by reducing selection bias, laboratory and operation-related procedures should be considered when determining whether a particular animal model is suitable.^35,36^ However, lost or euthanized animals do not necessarily reflect insufficient experimenter skills or procedural deficiencies within a laboratory facility. Indeed, such reputational concerns together with the perception that lost animals do not contribute to any relevant evidence hinder transparency, explaining in part the slow uptake and insufficient detail of reporting of animal attrition in the preclinical literature.^9,11,37^

Instead, we advocate for a perceptual shift towards describing the observable differences in characteristics of lost and retained animals. This can reveal valuable insights into limitations of inferences, particularly, when there is high attrition in the treatment but not the control group.^30,38^ The characteristics of lost animals within each experimental group should be systematically documented and published alongside the study results, which aligns with the ARRIVE guideline recommendations.^38^

While we emphasize the impact of collider stratification bias for *in vivo* research, *in vitro* research may also be susceptible to this type of selection bias due to exclusion of samples during the course of experiments.

Multidimensional assessments of animal welfare should include measurements of relevant causes of the animal welfare (such as initial infarct volumes). Explicitly reporting details about these variables and their association with the welfare score and the outcome could strengthen the knowledge about their role in the causal processes in the animal model and be informative to other researchers working with the same model.

Lastly, DAGs serve as a valuable tool to formulate *a priori* hypotheses about the underlying data generating processes. They visually communicate these hypotheses, aiding in the identification of sources of potential biases, and justify design and analysis choices.^1,20^ DAGs therefore lend themselves to the inclusion into pre-registration of preclinical experiments, especially to justify adjustment in cases when biases are suspected. They help facilitate a comprehensive understanding of the complex interplay between variables, thereby strengthening the interpretation of research findings and the robustness of the research design. In circumstances in which knowledge of the data generating process (i.e., DAG and functional relationships) is available, it may be possible to mitigate selection bias due to animal attrition in the statistical analysis phase.

## Conclusions

Even when conducting randomized, controlled, standardized laboratory animal experiments, researchers should be aware of the potential for selection biases that may pose a threat to internal validity. In our study, we have explored the mechanisms behind collider stratification bias, a specific type of selection bias. Though well-described in the field of epidemiology, we have demonstrated why collider stratification bias should be a concern in preclinical studies involving animal models, particularly those with severe side effects and high post-randomization attrition. We focused on scenarios in which the treatment had no actual effect on the outcome, but the researcher would have (falsely) observed one due to this type of selection bias. Collider stratification bias can strongly affect the conclusions drawn from such experiments, in turn influencing the direction of subsequent translational research. Our findings emphasize the importance of considering and addressing selection biases in preclinical research to improve the validity of study results, contributing to more accurate and reliable scientific discoveries.

## Authors’ contribution statement

TK and HGK conceived the study. AC, MP and JLR developed the study design. AC and HGK performed the literature review and designed the laboratory application. AC created the R code and performed the simulation study under the supervision of MP. AC prepared the figures. All authors interpreted the results. AC wrote the first draft of the manuscript. All authors critically reviewed the draft for important intellectual content and contributed to the final, revised version. JLR provided project supervision.

